# Development of Magnetic Particle Imaging (MPI) for Cell Tracking and Detection

**DOI:** 10.1101/2020.07.12.197780

**Authors:** Kierstin P Melo, Ashley V Makela, Natasha N Knier, Amanda M Hamilton, Paula J Foster

## Abstract

**Introduction:** Magnetic particle imaging (MPI) is a new imaging modality that sensitively and specifically detects superparamagnetic iron oxide nanoparticles (SPIONs) within a sample. SPION-based MRI cell tracking has very high sensitivity, but low specificity and quantification of iron labeled cells is difficult. MPI cell tracking could overcome these challenges.

**Methods:** MDM-AB-231BR cells labeled with MPIO, mice were intracardially injected with either 2.5 × 10^5^ or 5.0 × 10^5^ cells. MRI was performed in vivo the same day at 3T using a bSSFP sequence. After mice were imaged ex vivo with MPI. In a second experiment Mice received an intracardiac injection of either 2.5 × 10 ^5^ or 5 × 10 ^4^ MPIO-labeled 231BR cells. In a third experiment, mice were injected with 5 × 10 ^4^ 4T1BR cells, labelled with either MPIO or the SPION Vivotrax. MRI and MPI was performed in vivo.

**Results:** Signal from MPI and signal voids from MRI both showed more iron content in mice receiving an injection of 5.0 × 10^5^ cells than the 2.5 × 10^5^ injection. In the second experiment, Day 0 MRI showed signal voids and MPI signal was detected in all mouse brains. The MPI signal and iron content measured in the brains of mice that were injected with 2.5 × 10 ^5^ cells were approximately four times greater than in brains injected with 5 × 10 ^4^ cells. In the third experiment, in vivo MRI was able to detect signal voids in the brains of mice injected with Vivotrax and MPIO, although voids were fainter in Vivotrax labeled cells. In vivo MPI signal was only detectable in mice injected with MPIO-labeled cells.

**Conclusion:** This is the first example of the use of MPIO for cell tracking with MPI. With an intracardiac cell injection, approximately 15% of the injected cells are expected to arrest in the brain vasculature. For our lowest cell injection of 5.0 × 10^4^ cells this is ∼10000 cells.

## Introduction

Cellular MRI combines the ability to obtain high-resolution MRI data with the use of magnetic contrast agents for labeling specific cells, thereby enhancing their detectability.^1,2^ The most widely used cell labeling agents for cell tracking are magnetite (Fe_3_O_4_)-based superparamagnetic iron oxide (SPIO) nanoparticles. Commonly used iron oxide nanoparticles consist of a small (< 10 nm) iron oxide crystal core covered by a dextran coating bringing the total hydrodynamic size of the particles to approximately 20-50 nm (ultrasmall, USPIO) or 60-150 nm (standard, SPIO), respectively. Micron-sized iron oxide particles (MPIO) have also been used for preclinical cell tracking studies. MPIO are superparamagnetic microspheres with multiple small iron crystals (5-10nm) distributed throughout a polymer matrix that are relatively large (0.9 – 1.63 μm hydrodynamic size). Of the SPIO used for MRI cell tracking, MPIO have the highest iron content per particle, approximately 1pg Fe/particle which is roughly equivalent to 1.5 million standard SPIO particles or 4 million USPIO.^3^ The presence of SPIOs causes a distortion in the magnetic field and leads to signal hypo-intensities in iron-sensitive images (T2- and T2*-weighted images are most often used). Areas containing SPIO-labeled cells appear as regions of low signal intensity (signal voids) on MRI images, creating negative contrast. Many different cell types have been pre-labeled with iron particles and tracked with MRI, including mesenchymal stem cells,^4,5^ progenitor cells,^6^ dendritic cells,^7-9^ cancer cells^10-12^ and pancreatic islets^13^. This technique is highly sensitive, permitting the imaging of single cells *in vivo*^14,15^, under ideal conditions.

There are, however, several limitations of iron-based MRI cell tracking. The first is low specificity due to other low-signal regions in T2/T2* images; i.e. SPIO-labeled cells in the lung or in a region of hemorrhage cannot be detected. Although ultra-short echo time imaging methods have been developed for producing positive contrast from iron-labeled cells these too have similar problems with specificity.^16^ Second, quantification of iron-induced signal loss is complicated. Our group and others have shown that the degree of signal loss produced by SPIO-labeled cells is only linear at low iron concentrations.^17^ Typically, the degree of contrast (how black is it) or the volume of signal loss (how big a void is there) is measured from these images. Although some studies have used MR-based relaxometry to show a linear relationship between R2*values and SPIO-labeled cell concentration in samples^18^ the *in vivo* cell quantification by this method is more complicated. Overall, with iron-based MRI cell tracking there are issues with specificity and iron-labeled cell number cannot be accurately determined.

Magnetic Particle Imaging (MPI) is a new imaging modality that directly detects SPIOs.^19,20^ MPI cell tracking may address the limitations presented by SPIO-based cell tracking. First, the MPI signal is only generated when the magnetic moments of the SPIOs rotate; this change in magnetization is in response to the application of an excitation field and is localized only to the SPIO within a region devoid of a magnetic gradient. This results in positive “hot-spot” contrast that provides spatial localization without ambiguity. This is because there is no signal from within the subject as biological tissues neither generate nor attenuate MPI signals. Second, the MPI signal is linearly quantitative with iron concentration, and therefore the number of SPIO labeled cells can be directly calculated. The sensitivity of MPI derives from the direct detection of the electronic magnetization of SPIO, which is 10^8^ times larger than the nuclear magnetization of protons seen in MRI.^21^ This translates to a theoretical MPI sensitivity in the hundreds of cells with current hardware and available SPIO. The highest cell detection sensitivity to date was reported to be 250 cells *in vivo*.^22^

The ideal SPIOs for MPI are still not known. Both MPI sensitivity and resolution are closely related to the physical properties of a SPIO. The resolution of MPI is driven primarily by the interaction of the nanoparticle and the magnetic field gradient. Theoretical modeling predicts that resolution improves with increasing core size.^23^ However, this has not always been observed experimentally. The sensitivity of MPI depends on both nanoparticle and scanner specific factors. Nanoparticle factors include the strength of the nanoparticle magnetization (the greater the magnetization the greater the MPI signal and therefore higher sensitivity) and the efficiency of the nanoparticle cell labeling (more iron per cell leads to higher sensitivity).

Currently the most commonly used SPIO for MPI has been a commercially available agent, Vivotrax from Magnetic Insight Inc. (USA). VivoTrax is a Ferucarbotran with multi-core/aggregated particles and coated with carboxy-dextran. Vivotrax has been used in MPI studies of mice to detect mesenchymal stem cells^24,25^, neural stem cells^26^, neural progenitor cells^27^, pancreatic islets^28^, T-cells^29^, and macrophages.^30,31^

Our lab has previously used MPIO to label metastatic cancer cells for detection in the mouse brain by MRI. In our previous studies MPIO-labeled cancer cells were administered via intra-cardiac injection which results in their arrest throughout the brain vasculature as individual cells or clusters of small numbers of cells. The high cellular iron loading created when labeling cells with MPIO (30-40 pg of iron/cell) permitted the detection of single cells in vivo by MRI.^10^ At this time there are no published reports of the use of MPIO-labeled cells for their in vivo detection by MPI. Therefore, the goal of this study was to evaluate if MPIO can be used for in vivo detection and quantification of cancer cells distributed in the mouse brain by MPI.

## Methods

### Iron Oxide Nanoparticles and Relaxometry

Two types of iron oxide nanoparticles were used in these studies: (i) Vivotrax (Magnetic Insight Inc, Alameda, CA, USA) and (ii) MPIO which have a diameter of 0.9 um (BangsLaboratories Inc, Fishers, IN). The particle relaxometer module (RELAX™) on the MOMENTUM MPI system (Magnetic Insight Inc., Alameda, California) was used to characterize MPIO and Vivotrax (2.8 ug Fe in 1 ul for each). In this mode, the localizer gradient field is switched off and a negative magnetic field is turned on and then switched to a positive field (and vice versa). As a result, iron nanoparticles are driven from a negative magnetic saturation to positive (positive scan) and vice versa (negative scan). This measures the point spread function (PSF) of the nanoparticles and allows for measurements of the MPI signal per iron content (sensitivity) and full-width half-maximum (FWHM; spatial resolution). A narrower tracer response indicates superior spatial resolution and a greater signal intensity per unit iron indicates superior sensitivity.

### Calibration Line Preparation

Calibration lines were generated for use in quantifying iron content in brains after iron-labeled cell injection. To construct each line, samples of each nanoparticle were scanned in the same mode as images being analyzed (In vivo = 3D isotropic, ex vivo = 2D default). Samples of either Vivotrax or MPIO were diluted in PBS into 100%, 75%, 50%, 37.5%, 25%, 10%, 7.5%, 5%, 3.75%, 2.5% aliquots. 1ul of each dilution was then pipetted into capillary tubing and spaced out 2cm from each other on the MPI sample bed to be imaged, 5 samples at a time. The field of view (FOV) was 12cm × 6cm × 6cm.

### Cell Culture and Labeling

Two cancer cell types were used in these studies: (i) human MDA-231-BR brain metastatic breast cancer cell line and (ii) murine 4T1BR5 metastatic breast cancer cell line. Both cell lines were maintained at 37°C and 5% CO_2_ in DMEM (Gibco, Thermo Fisher Scientific, Inc., Waltham, MA, USA) supplemented with 10% fetal bovine serum and 1% penicillin streptomycin antibiotic. Cells were passaged every 2-3 days. For cell labeling with MPIO, adherent cells were incubated with 25 μg Fe/mL MPIO beads for 24 hours. Cells were washed three times with Hanks balanced salt solution (HBSS) and then trypsinized with 0.25% Trypsin-EDTA. The cells were then collected and washed another three times with HBSS to remove unincorporated MPIO before cell injection and *in vitro* evaluation. For cell labeling with Vivotrax ™, cells were grown for 2-4 days until they reach 80-90% confluency. A labeling mixture was prepared using 2 falcon tubes of 2.5 mL of serum free low glucose DMEM. In tube A, 60 μL of stock protamine sulfate was added and vortexed to mix. In tube B, 90 μL of VivoTrax was added and vortexed to mix. 20uL of stock heparin was then added to tube B and then vortexed to mix. Old media was removed from cells and rinse once with HBSS. Tubes A and B were combined and vortexed. This labeling mix was added to the flask and incubated at 37°C, 5% CO2 for 2-4 hours. 5 mL of complete low glucose DMEM was added to the flask after the 2-4 hours and incubated at 37°C, 5% CO2 overnight. On day 2 cells were washed 3 times with HBSS and then trypsinized with 0.25% Trypsin-EDTA. The cells were then collected and washed another three times with HBSS to remove unincorporated Vivotrax™ before cell injection and *in vitro* evaluation. This approach for cell labeling is what we routinely do for labeling cells with ferumoxytol nanoparticles (ie. Feraheme) and was used in this study for cell labeling with Vivotrax because cells could not be labeled efficiently with simple co-incubation, as they could be with MPIO. In both scenarios, cell viability after labeling was assessed using the trypan blue exclusion assay and labeling efficiency was assessed by Perl’s Prussian blue (PPB) staining by counting the number of positively stained cells in 3 random 40x magnification fields under the microscope.

### Mouse Model

Female nude (nu/nu) (6-8 weeks; Charles River Canada or USA) or NSG (NOD/SCID/ILIIrg^−/−^) mice were obtained and cared for in accordance with the standards of the Canadian Council on Animal Care, under an approved protocol by the Animal Use Subcommittee of Western University’s Council on Animal Care and Use Committee. Mice were anesthetized with isoflurane administered at 2% in oxygen and iron-labeled cancer cells were injected intracardially into the left ventricle of the heart under ultrasound guidance using a Vevo 2100 ultrasound system (Visual Sonics Inc.). The delivery of cells to the brain after intracardiac injection is related to the cardiac output, with only 15-20% of cardiac output reaching the brain. Based on this, we estimate that a similar percentage of iron-labeled cells will be initially delivered to the brain with a technically accurate injection. In our previous studies which have used intracardiac injection of iron-labeled cells to study brain metastasis, MRI is performed on the same day of the injection to verify delivery of cells to the brain. We have demonstrated that the number of signal voids, caused by iron-labeled cells, increases with the number of cells injected. When considering cell detection sensitivity for MPI it is worth noting that cells injected this way will be distributed throughout the entire brain, not in a similar region all together.

Experiment 1 (*Ex vivo* MPI): Mice received an intracardiac injection of either 2.5 × 10^5^ (n=2, ∼50,000 cells in brain) or 5 × 10^5^ (n=2, ∼100,000 cells in brain) MPIO-labeled 231BR cells in 0.1 mL of HBSS. One mouse from each group was imaged with MRI *in vivo* (as described below) on the day of the injection (day 0) and then the mice were euthanized and the fixed mouse heads were shipped to Michigan State University (East Lansing, MI) to be imaged with MPI on a Momentum MPI system by co-author AM. At this time, we did not have an MPI system installed at Robarts Research Institute. The goal of this preliminary experiment was to determine if MPIO-labeled cells could be detected in this mouse brain model system. MPI data was acquired with default 2D scans using a 6 × 6 × 4 cm FOV and a 5.7 T/m gradient.

Experiment 2 (*In vivo* MPI): Mice received an intracardiac injection of either 2.5 × 10^5^ (n=3, ∼50,000 cells in brain) or 5 × 10^4^ (n=3, ∼10,000 cells in brain) MPIO-labeled 231BR cells in 0.1 mL of HBSS. These mice were imaged *in vivo* on the day of the injection (day 0) with MRI and then MPI on a Momentum MPI system at Robarts Research Institute. 3D images were acquired using a 3 T/m gradient, 35 projections and a FOV 12 × 6 × 6 cm, with a total scan time ∼1 hour per mouse (the 3D high sensitivity mode on the Momentum system).

Experiment 3: Mice received an intracardiac injection of 5 × 10^4^ Vivotrax- or MPIO-labeled 4T1BR5 cells (n=4 per group, ∼10,000 cells in brain). These mice were imaged *in vivo* on the day of the injection (day 0) with MRI and MPI, as in Experiment 2.

MPI images were analyzed utilizing Horos imaging software. (Horos is a free and open source code software program that is distributed free of charge under the LGPL license at Horosproject.org and sponsored by Nimble Co LLC d/b/a Purview in Annapolis, MD USA). Images were displayed in full dynamic range and total MPI signal was calculated. Areas of interest from 3D images were manually outlined, slice by slice, creating a 3D volume. In 2D images, the areas of interest were manually outlined in a single slice. The mean signal from these ROI’s were then multiplied by the ROI volume/area to determine the total MPI signal. For the nanoparticle samples the total MPI signal was plotted against iron content to derive the calibration lines. This relationship was used to quantify iron content (x) in mouse brains. Where the total MPI signal was substituted for y, (m) is the slope of the line and (b) is the y-intercept in y= m × +b. All MPI images were analyzed in the same way to ensure consistency.

### Magnetic Resonance Imaging

*In vivo* proton (^1^H) MRI for all mice was performed at 3 Tesla using a GE MR750 system equipped with an insertable gradient coil and a solenoidal mouse head radiofrequency coil. While imaging, mice were anesthetized with 2% isoflurane in oxygen. A 3D balanced steady state free precession pulse sequence was used. Image resolution was 200 × 200 × 200 μm and sequence parameters were as follows: FOV = 50 × 50 mm with a half phase FOV, matrix = 250 × 250, flip angle = 35°, receiver bandwidth = +/−41.67 kHz, repetition time/echo time (TR/TE) = 4.2/2.1 ms, 2 signal averages and 8 phase cycles resulting in a scan time of ∼30 minutes.

MRI data was also visualized and analyzed using Horos imaging software. Brain images were assessed for the presence of signal voids attributable to iron-labeled cells arrested in the mouse brain vasculature after the intracardiac injection. Iron-labeled cells in the brain were quantified by determining the percentage of black pixels by drawing a region of interest (ROI) around the whole brain and setting a threshold value based on the mean signal intensity value of a signal void ± 2 standard deviations. The total number of black pixels below this threshold value was obtained from the entire brain volume signal intensity histogram. The number of black pixels was divided by total number of pixels to calculate the percentage of black pixels.

### Statistical Analysis

Data are presented as mean +/− standard deviation. Linear correlations were conducted between total MPI signal and iron content. Linear regression was calculated to create equations used for iron content quantification and the goodness of fit R^2^ value (Pearson’s correlation coefficient). These analyses were conducted using Prism software (8.0.2, GraphPad Inc.), where p< 0.05 was considered statistically significant.

## Results

### Evaluation of Vivotrax and MPIO performance by Relaxometry and MPI

The relaxometer mode on the MPI system was used to compare the performance of MPIO to Vivotrax. Figure 1A shows the spectra for both iron nanoparticles. The amplitude of the MPI signal for MPIO was slightly higher than Vivotrax; the relative sensitivity was 1.23 for MPIO versus 1.0 for Vivotrax. Figure 1B compares the resolution of the particles using the normalized MPI signal. The resolution of MPIO was almost twice that of Vivotrax; 3.0 versus 1.65 mm for a 6.1 T/m gradient. An example of the images of samples measured to generate calibration curves are shown in Figure 1C for MPIO and the calibration line generated from this data is shown in 1D. There was a strong linear relationship between iron content and MPI signal (arbitrary units, A.U.) for both MPIO (R^2^ = 0.974, P < .001) and Vivotrax (R^2^ = 0.9678, P < .001). The equation of the line was: MPI Signal = 0.0426 (Iron Content) + 0.179 for MPIO and MPI Signal =0.0017 (Iron Content) + 0.1706 for Vivotrax. Using this relationship, iron content could be determined for a given MPI signal.

**Figure 1:**
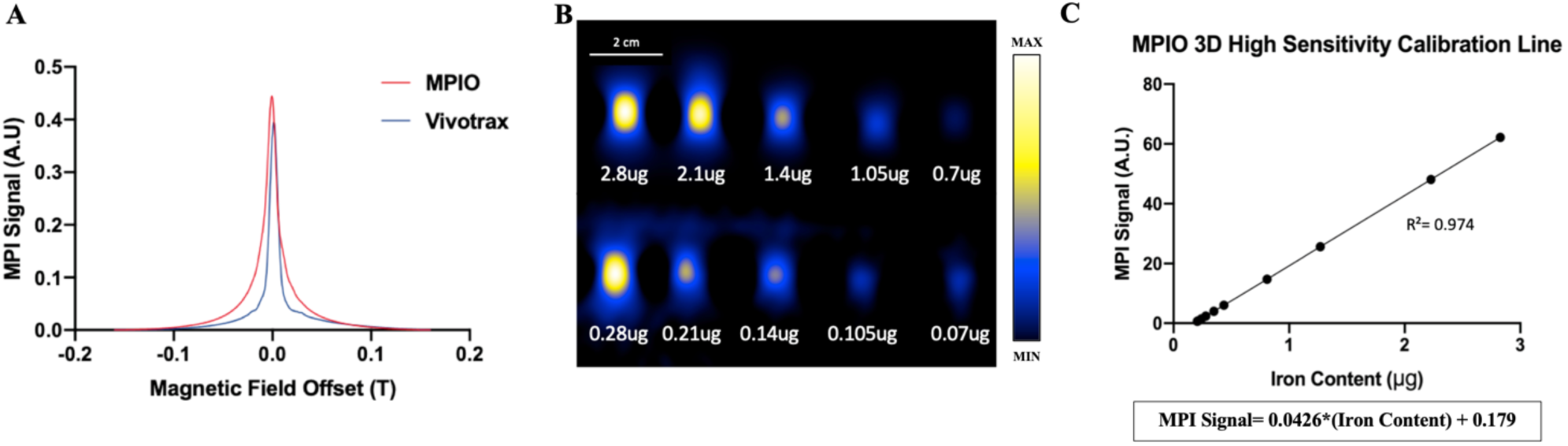
(A) Relaxometer data that compares the signal amplitudes of both MPIO and Vivotrax nanoparticles to show expected sensitivity as well as resolution by the FWHM. (B) Diluted samples of MPIO used to create a calibration line. (C) The relationship between iron content and MPI signal is linear at both high and low concentrations of iron.

### Imaging

PPB staining confirmed the successful labeling of cancer cells with either MPIO or Vivotrax. Labeling efficiency was greater than 90% for MPIO and 70% for Vivotrax. Labeling with either nanoparticle did not change cell viability; > 95% before and after labeling. Figure 2 shows representative *in vivo* MRI and *ex vivo* MPI images from Experiment 1 for mice injected with either 2.5 × 10^5^ (Fig 2 b & d; n=2) or 5 × 10^5^ (Fig 2 a & c; n=2) MPIO-labeled 231BR cells. One mouse from each group was imaged *in vivo* with MRI. MR images showed the characteristic signal voids throughout the brain (Fig 2 a & b) representative of iron-labeled cells arrested in the brain on the day of the intra-cardiac injection, consistent with numerous previous MRI studies conducted in our lab. The signal voids were quantified by measuring the percentage of black pixels in the whole brain. These values were 1.04% for the 2.5 × 10^5^ cell injection and 2.77% for the 5 × 10^5^ cell injection. MPIO-labeled 231BR cells were detected by MPI in all fixed brains (Fig 2 c & d). Quantification of the MRI and MPI data is shown in Table 1. This preliminary experiment did not use sufficient animal numbers to perform a statistical analysis of this data but did demonstrated that MPIO-labeled cells arrested in the mouse brain could be detected and quantified by MPI. In Experiment 2 MPI was performed *in vivo* and fewer cells were injected to learn more about MPI cellular detection sensitivity.

**Table 1:**
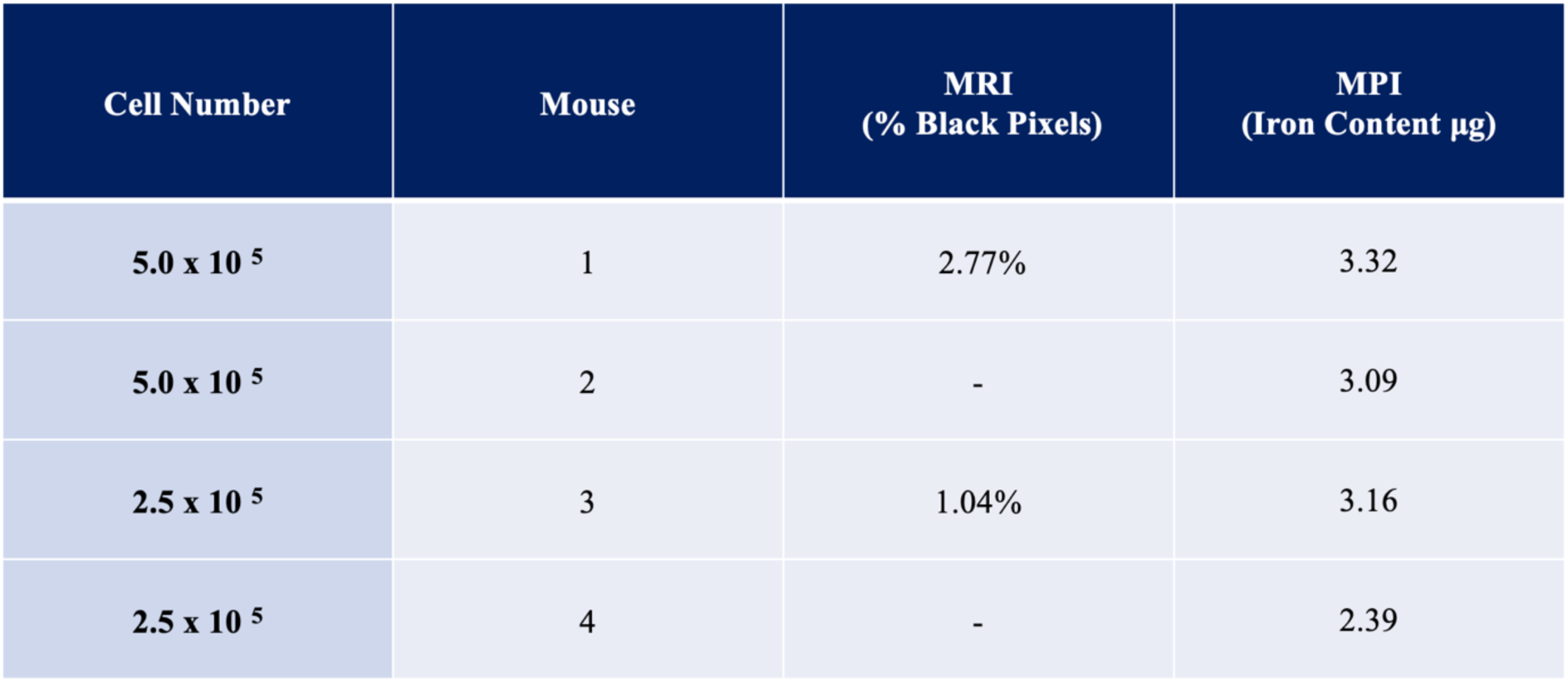
Summary of experiment 1 comparing cell number to its corresponding % black pixel from MRI images and iron content calculated from MPI signal.

**Figure 2:**
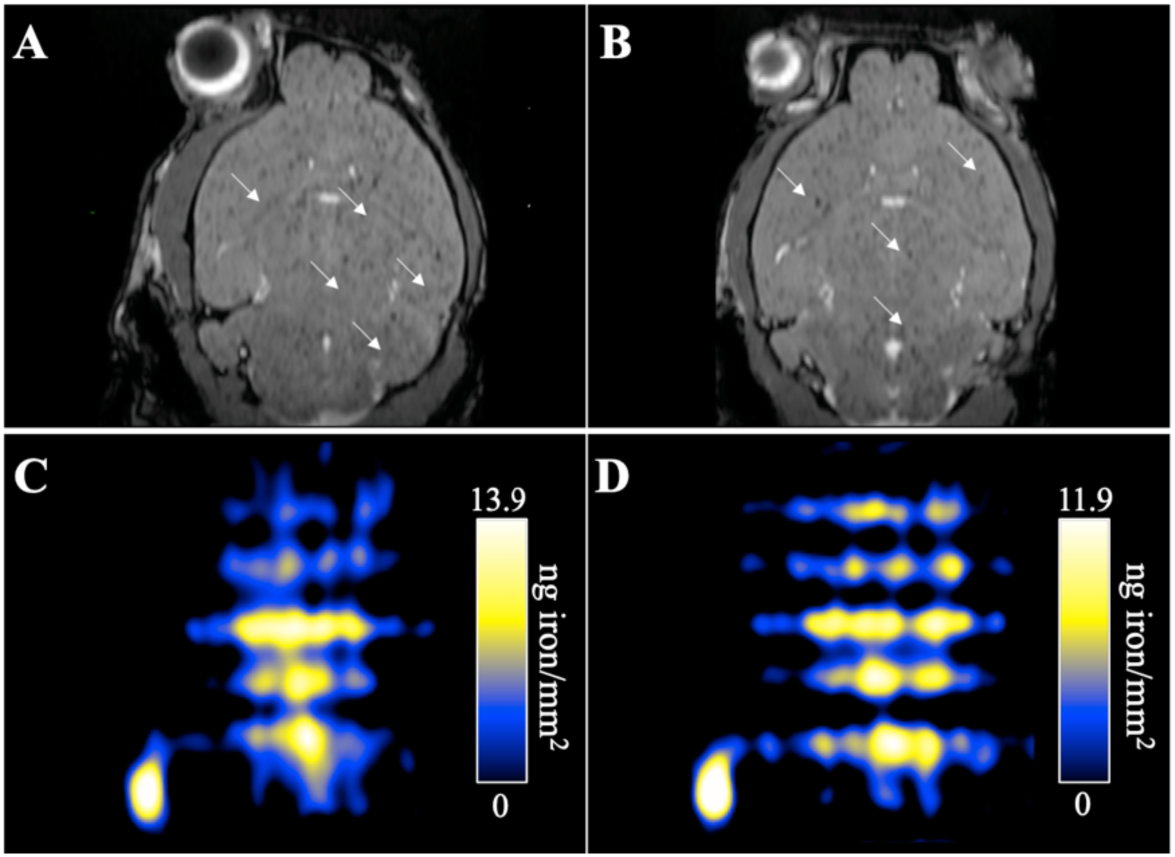
(A) MRI of mouse brains injected with 5.0 × 10 ^5^ (B) and 2.5 × 10 ^5^ 231BR cells labeled with MPIO. (C) MPI of mouse brain injected with 5.0 × 10 ^5^ (D) and 2.5 × 10 ^5^ 231BR cells labeled with MPIO.

In Experiment 2, mice were injected with either 2.5 × 10^5^ (n=3) or 5 × 10^4^ (n=3) MPIO-labeled 231BR cells and were imaged *in vivo* with MRI and MPI. Day 0 MRI showed signal voids and MPI signal was detected in all mouse brains. The MPI signal and iron content measured in the brains of mice that were injected with 2.5 × 10^5^ cells was approximately four times greater than in brains injected with 5 × 10^4^ cells (average values 1.4 vs 0.37 ug).

In Experiment 3 mice were injected with 5 × 10^4^ MPIO-labeled 4T1BR5 cells (n=4) or 5 × 10^4^ Vivotrax-labeled 4T1BR5 cells (n=4) and imaged *in vivo* with MRI and MPI. Day 0 MRI showed distinct signal voids in mouse brains injected with MPIO labeled cells (Figure 4A), consistent with previous experiments. MPI signal was detected in all mice injected with 5 × 10^4^ MPIO-labeled cells, a representative image is shown in Figure 4C. The MPI signal in these images appear smoother as a result of the scan mode used for in vivo MPI which trades some resolution for sensitivity. The iron content measured from MPI of the brains of mice which were injected with 5 × 10^4^ MPIO-labeled 4T1BR5 cells was not significantly different from that from Experiment 2 mice which were injected with 5 × 10^4^ MPIO-labeled 231BR cells; average brain iron content was 0.28 ug for 4T1BR5 cells and 0.37 ug for 231BR cells. Vivotrax-labeled cells appeared as very faint signal voids in MRI which were more difficult to detect and not distinct enough to permit quantification (Figure 4B). No MPI signal was detected in the brains of mice injected with 5 × 10^4^ Vivotrax-labeled 4T1BR5 cells.

**Figure 4:**
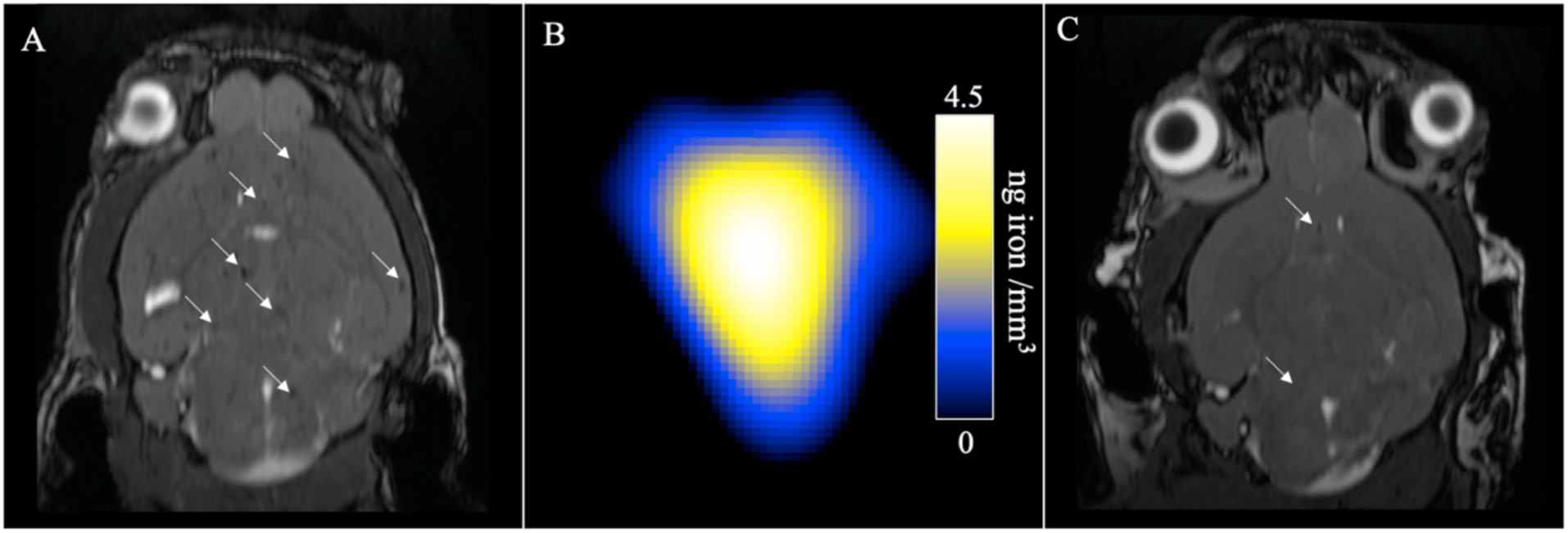
(A) MRI of mouse brain injected with 5.0 × 10 ^4^ 4T1BR5 labeled with MPIO. (B) MPI scan (C) MRI of mouse brain injected with 5.0 × 10 ^4^ 4T1BR5 labeled with Vivotrax ™.

## Discussion

This is the first study to demonstrate that MPIO-labeled cells can be detected and quantified *in vivo* by MPI. MPI spatial resolution and sensitivity are heavily influenced by the physical properties of the nanoparticle, such as the effective core size, relaxation time and size distribution. One way to improve resolution is to increase the (effective) magnetic core size of the particles. The Langevin model of superparamagnetism predicts a cubic improvement of resolution with increasing core size. However, the temporal response, or magnetic relaxation, of particles increases with core size and this leads to blurring which hinders the improvements in resolution. Here, magnetic relaxation can be thought of as the non-instantaneous response of the ensemble magnetization to the applied field.^32^ Tay *et al.* have found that improved resolution with increasing magnetic core size follows the predictions up to approximately 25 nm when the effects of SPIO rotational times become significant.^20^ Sensitivity depends, among other factors, on the strength of the nanoparticle magnetization. The magnetization can be increased by enlarging the iron core diameter, as the strength of the MPI signal increases by the third power of the iron core diameter.^33^

In the early days of MPI, commercially available SPIOs used for MRI were evaluated and Resovist (which is the same as Vivotrax) showed a good MPI performance. Although widely used, Vivotrax is now not considered optimal for MPI because it has a bimodal size distribution, predominantly containing small cores ∼ 5 nm in diameter with a small fraction (30%) of multi-core aggregates with an effective size of 24 nm.^34^ The individual cores are too small to magnetize significantly and so the MPI signal only comes from the clustered multi-core structures. One approach being taken to improve MPI sensitivity is to design similar particles but with a bigger fraction of the aggregates. Eberbeck et al. have designed Nanomag-MIP particles which are similar to Vivotrax, they are composed of individual cores between 3-8 nm along with multi-core aggregates with an effective core size of 19 nm.^35^ However, for Nanomag-MIP 80% of the nanoparticles are of the larger size and the MPI signal is two times larger than that of Resovist Yoshida et al. showed that fractionation of Resovist can improve the MPI signal by 2.5 times.^36^ Magnetic Insight Inc. have recently made available for purchase Vivotrax *Plus*™, which is size-filtered Vivotrax, to generate a larger fraction of clustered particles with an ideal size, and is said to have three times the signal and improved resolution compared to Vivotrax.^37^ Another approach is the synthesis of homogeneously distributed single-core SPIOs with a dedicated iron core diameter for ideal MPI characteristics. Ferguson et al. observed increasing MPI signal with increasing magnetic core size (14-27 nm) for monodisperse single core particles and up to three times greater signal intensity per unit iron compared to multi-core Vivotrax.^34^

MPIO are quite different from the nanoparticles typically used for MPI. MPIO consist of multiple small cores (∼5-10nm)^35^ embedded in polystyrene matrix. They have a very high iron content (63%) and because of the clustered nature of the iron cores they can be regarded as having one very large superparamagnetic core. The 0.9 μm sized MPIO we have used have a broad size distribution, the specified range is 0.5-2 μm, however all of these particles would be expected to contribute to the MPI signal.

Figure 1 shows that the MPI signal generated by the MPIO has reduced resolution compared to Vivotrax, consistent with theory. The low resolution is apparent in MPI images of MPIO-labeled cells in the mouse brain which show one large area of signal. Unlike, with MRI, it is not possible to determine where within the brain the cells are located from the MPI images. Figure 1 shows that the MPI signal for MPIO is greater than that for Vivotrax for equal iron concentrations. This suggests that the net change in magnetization of the MPIO is greater than the net magnetization change of the same mass of Vivotrax, since the MPI signal results from the change in magnetization of an excited particle. The high MPI signal produced by MPIO and the high intracellular iron content with MPIO labelling both contribute to our increased ability to detect MPIO-labelled cells distributed throughout the mouse brain.

We were able to detect and measure MPI signal in the brains of mice injected with as few as 5.0 × 10^4^ MPIO-labeled cells; an intra-cardiac injection of 5 × 10^4^ cells is estimated to deliver 10,000 cells to the mouse brain. We did not detect MPI signal in the brain when 5.0 × 10^4^ Vivotrax-lableled cells were injected. This is, in part, due to the fact that cell labeling with MPIO is more efficient than for Vivotrax (∼90% versus 70% of cells labeled) and the amount of iron per cell is significantly greater for MPIO than Vivotrax labeling (30-40 pg/cell versus 10-15 pg/cell). Our MRI data for Vivotrax cells in the brain supports this; signal voids were considerable fainter and harder to detect in MR images of mouse brains which received Vivotrax-labeled cells. Furthermore, the MPI signal generated from MPIO is higher than Vivotrax (Figure 1). While Vivotrax is currently the most common SPIO used for MPI, our results show that MPIO-labeled cells can be detected and quantified in this mouse brain model more readily than Vivotrax-labeled cells. It is worth noting that MPIO are inert, nonbiodegradable particles only suitable for preclinical experimental studies.

MPI provided information which was not attainable with MRI. With knowledge of the amount of iron per cell achieved with labeling the cell number could be estimated. Our routine measurements of mean iron/cell after labeling the cancer cell lines used in this study with MPIO are consistently in the range of 30-40 pg of iron/cell, using inductively-coupled mass spectrometry (ICP-MS). As described above, with an estimate of 20% of injected cells arresting in the brain the intra-cardiac injections of 5.0 × 10^5^, 2.5 × 10^5^ and 5 × 10^4^ cells will deliver approximately 100,000, 50,000 and 10,000 cells to the brain, respectively. If we use the lower value (30 pg) for MPIO labeling of cells, for example, this amounts to approximately 3.0, 1.5 and 0.3 ug of iron in the brain due to iron-labeled cell burden. As shown in Table 2, our data falls within these approximations.

**Table 2:** Summary of all experiments, showing cell number injected, estimated number of arrested cells in the brain, estimated amount of iron per cell and estimated amount of iron in the brain compared to actual measured iron content in the brain from MPI images.

| Number of Cells Injected (IC) | Estimate of Number of Cells Arrested in Brain | Estimate of Iron per cell (pg) | Estimate of Iron in the Brain (μg) | MPI Measurement of iron in brain (μg) |
| --- | --- | --- | --- | --- |
| $5.0 \times 10^4$ | 10 000 | 30 pg | 0.3 μg | 0.2 - 0.7 μg |
| $2.5 \times 10^5$ | 50 000 | 30 pg | 1.5 μg | 1.3 – 3.2 μg |
| $5.0 \times 10^5$ | 100 000 | 30 pg | 3 μg | 3.1 - 3.3 μg |

There are a number of factors which make it impossible to calculate true cell number in our study. First, the value for iron/cell that we measure from ICP-MS or MPI is an average value, some cells will contain more iron, some less. Second, the number of cells delivered to the brain by intra-cardiac injection is an estimation. We used an estimate of 20% of cardiac output to the brain resulting in 20% of cells delivered; published values are between 15-20%^36^. Intra-cardiac injections are also technically challenging and, while we have significant expertise, not every injection is likely to deliver the exact same number of cells. Lastly, there may be an upper limit to the number of cells that arrest and then persist in the brain. For these last two reasons, there are unlikely to be double the number of cells arrested in the brain with double the number injected. Considering all these caveats our values for iron content measured by MPI are in good agreement with the estimate of iron expected in the brain. In conclusion, we have demonstrated that MPIO-labeled cells can be detected in vivo in a model where cells are dispersed throughout the mouse brain.

## Notes

### Competing Interest Statement

The authors have declared no competing interest.

### Summary of Updates

Natasha N Knier added as a co-author.

